# SNPPar: identifying convergent evolution and other homoplasies from microbial whole-genome alignments

**DOI:** 10.1101/2020.07.08.194480

**Authors:** David J. Edwards, Sebastián Duchêne, Bernard Pope, Kathryn E. Holt

**Author notes:** Correspondence: David Edwards.

## Abstract

Homoplasic single nucleotide polymorphisms (SNPs) are considered important signatures of strong (positive) selective pressure, and hence of adaptive evolution for clinically relevant traits such as antibiotic resistance and virulence. Here we present a new tool, SNPPar, for efficient detection and analysis of homoplasic SNPs from large WGS datasets (>1,000 isolates and/or >100,000 SNPs). SNPPar takes as input a SNP alignment, tree and annotated reference genome, and uses a combination of simple monophyly tests and ancestral state reconstruction (ASR, via TreeTime) to assign mutation events to branches and identify homoplasies. Mutations are annotated at the level of codon and gene, to facilitate analysis of convergent evolution.

Testing on simulated data (120 *Mycobacterium tuberculosis* alignments representing local and global samples) showed SNPPar can detect homoplasic SNPs with very high sensitivity (zero false-positives in all tests) and high specificity (zero false-negatives in 89% of tests). SNPPar analysis of three empirically sampled datasets (*E. anophelis, B. dolosa* and *M. tuberculosis*) produced results that were in concordance with previous studies, in terms of both individual homoplasies and evidence of convergence at the codon and gene levels. SNPPar analysis of a simulated alignment of ∼64,000 genome-wide SNPs from 2000 *M. tuberculosis* genomes took ∼23 minutes and ∼2.6 GB of RAM to generate complete annotated results on a laptop. This analysis required ASR be conducted for only 1.25% of SNPs, and the ASR step took ∼23 seconds and 0.4 GB RAM.

SNPPar automates the detection and annotation of homoplasic SNPs efficiently and accurately from large SNP alignments. As demonstrated by the examples included here, this information can be readily used to explore the role of homoplasy in parallel and/or convergent evolution at the level of nucleotide, codon and/or gene.

**Impact statement:** DNA sequences of bacterial pathogens are mutating all the time; most changes are deleterious or neutral, but sometimes a mutation leads to functional change that allows the pathogen to evade a potential threat. These random mutational changes (single nucleotide polymorphisms, or SNPs) are so very rarely beneficial, that when they do arise in parallel in distantly related isolates (known as homoplasic SNPs) this indicates that the change may be positively selected because it confers an adaptive advantage to the bacteria.

Finding homoplasic SNPs in large sets of bacterial genomes is challenging as current tools require substantial time and computational resources to run. Here we present SNPPar, a software program to efficiently and accurately automate the detection and annotation of homoplasic SNPs from large whole-genome sequence data sets. We use simulated data to demonstrate accuracy of the program, and re-analyse published datasets using SNPPar to illustrate how the results can be used to gain insights into the evolution of antibiotic resistance and other traits.

We envisage SNPPar will help facilitate the undertaking of long-term, real-time surveillance of bacterial pathogens, and their adaptive evolutionary response to interventions and control measures such as new drugs or vaccines.

**Data summary:** The authors confirm all supporting data, code and protocols have been provided within the article, through supplementary data files or other online sources as indicated in the article.

New content generated for this paper is:

1. SNPPar code is available from https://github.com/d-j-e/SNPPar. The version described here is v1.0.
2. A GitHub repository containing the full protocol, ‘in-house’ code and data used to carry out the validation and performance testing is available at https://github.com/d-j-e/SNPPar_test. This repository includes all the simulated and real data sets used here.

**Data statement:** The authors confirm all supporting data, code and protocols have been provided within the article, through supplementary data files or other online sources as indicated in the article.

## 1. Introduction

Bacterial pathogen populations are under strong selection from antimicrobials and host immune defences, and there is increasing interest in utilising whole genome sequencing (WGS) data to detect adaptive evolution in response to these strong selective pressures. Of particular interest in the field are repeated or convergent evolutionary patterns^1,2^ in response to specific selective pressures (such as drug exposure, or within-host evolution), which imply predictability of adaptive responses that could be used to track disease progression, and to guide selection of vaccine and drug targets or therapeutic strategies that are more difficult for bacteria to evade through adaptive evolution^3,4^.

A key signature of adaptive evolution is the presence of homoplasies in the population^2,3,4^. Homoplasic traits are those that have been gained (or lost) independently in two or more lineages since their divergence from a common ancestor, in contrast to those traits that were gained or lost only once in a population and are shared by virtue of vertical inheritance from a common ancestor^5^ (see **Figure 1**). Extending this to single nucleotide polymorphisms (SNPs), homoplasic SNPs are those where the same derived nucleotide is present in two or more lineages due to independent mutation events that occurred since their divergence from a common ancestor (which harboured a distinct ancestral nucleotide). Under the infinite sites model of molecular evolution^6^, the same substitution event should not be observed multiple times in the absence of positive selection, thus homoplasic SNPs are considered important signatures of adaptive evolution.

**Figure 1.**
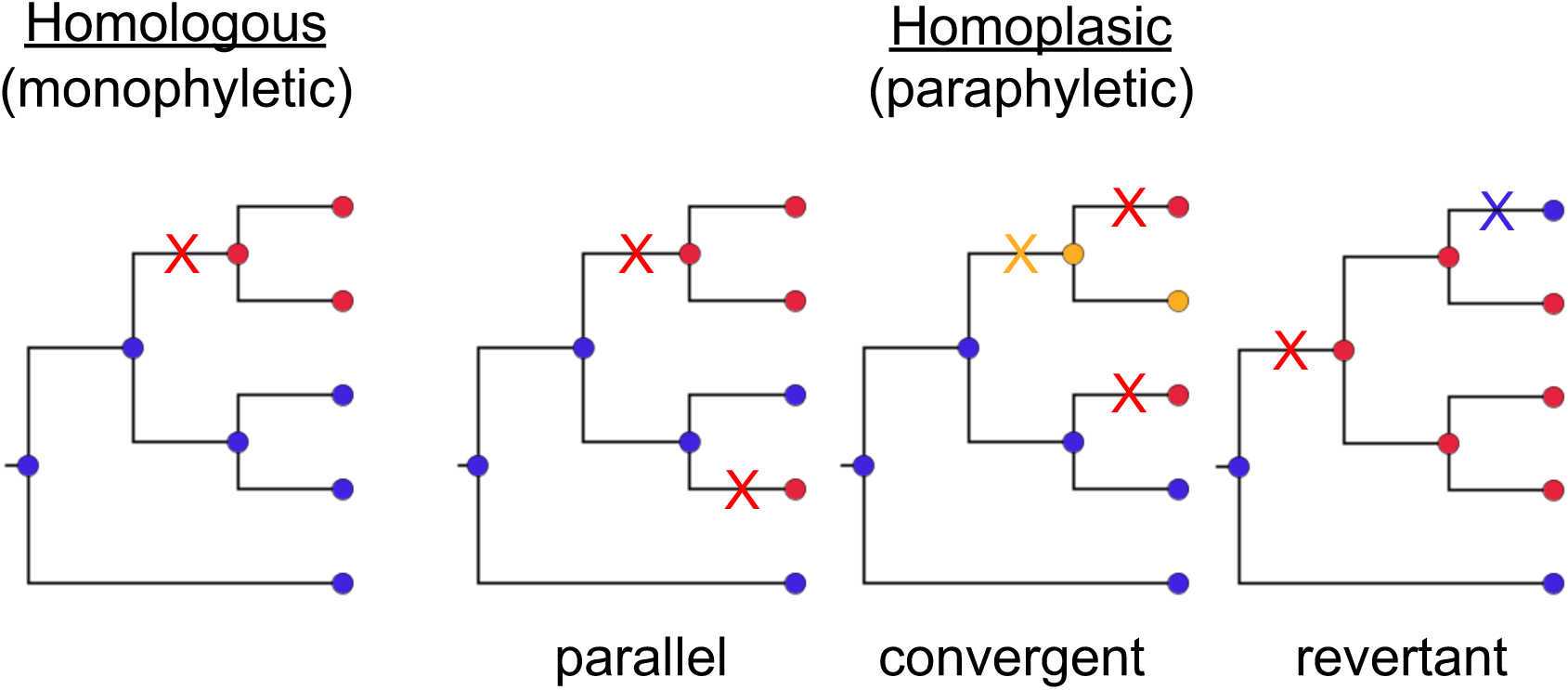
Example trees demonstrating the relationship between homology (monophyly) and homoplasy (paraphyly). Homoplasic SNPs can be the result of parallel, convergent or revertant mutation events. Different colours of nodes indicate different nucleotide bases (*e*.*g*. blue, adenine (A); red, thymine (T); yellow; guanine (G)). Different coloured crosses indicate mutations giving rise to the SNPs.

Homoplasic SNPs can arise through different series of mutation events, classed as parallel, convergent or revertant evolution (see **Figure 1**). Parallel homoplasic SNPs are those where the same substitution, *e*.*g*. A to T, occurs independently at the same site in multiple diverged lineages; convergent homoplasic SNPs are those where the same nucleotide arises in diverged lineages through a distinct series of substitution events at the same site (*e*.*g*. A to T in one lineage, and A to G to T in another lineage; see **Figure 1**). Revertance refers to restoration of a derived nucleotide back to the ancestral nucleotide, resulting in sharing of the ancestral nucleotide through mechanisms other than direct inheritance (*e*.*g*. a prior mutation of A to T is reversed in a descendant subpopulation by a subsequent substitution of T to A; see **Figure 1**). Note that because selection acts on the function of encoded proteins, convergence can also be examined at the level of codons, whereby independent DNA mutations (*e*.*g*. a third or fourth allele at the same SNP site, or a SNP at another site in the same codon) result in the same or functionally-equivalent amino acid substitution; or at the level of genes, whereby independent mutations affecting different codons have similar functional effects on the encoded protein’s structure and function.

Homoplasic SNPs can be produced by spontaneous mutation (*i*.*e*. errors during chromosome duplication prior to cell division) but can also arise via recombination. Whilst recombination is a major driving force of evolution in many bacterial populations, in clonal pathogens that exhibit very little recombination, most new variation is acquired through spontaneous mutations. Genomics-based studies aiming to understand clonal evolution and positive selection typically seek to identify and remove homoplasic SNPs that result from recombination^7,8^, which improves the resolution of vertical patterns of evolution in a phylogenetic tree^9^ and preserves homoplasic SNPs that arose through spontaneous mutation to be examined as signals of positive selection across the genome.

A general approach to discovering homoplasic SNPs from whole genome alignments is to construct a phylogenetic tree from the alignment of all SNP sites, then probabilistically infer ancestral states for each SNP site at internal nodes of the tree using ancestral state reconstruction (ASR).^10^ Example implementations of maximum likelihood (ML) approaches to ASR include tools developed before WGS such as FastML^11,12,13^ or PAML^14^, and recently more efficient tools developed for genome-scale datasets such as TreeTime^15^. These ASR tools typically report inferred internal node sequences, thus further analysis is required to infer mutation events and localise these to branches of the tree; to identify homoplasic SNPs; and to distinguish between parallel, convergent and revertant homoplasic SNPs. Further analyses are also required to predict coding effects of SNPs in order to identify convergence at the amino-acid or gene level. Currently, at least two programs specifically target the discovery of homoplasic SNPs from microbial whole-genome alignments: HomoplasyFinder^16^ and TreeTime^15^. Whilst both find homoplasic sites, the outputs are limited and require substantial further processing for most applications. HomoplasyFinder takes as input a tree and a SNP alignment, and reports sites with a consistency index <1 as homoplasic^16^. However, it does not identify the type of homoplasy, report details of the specific mutations and where they occur on the tree, nor include any functions for analysing potential coding consequences to detect convergence at codon level. TreeTime^15^ is developed primarily for inferring ML dated phylogenies and conducting ASR, and its ASR function is much faster than FastML or PAML (increase in execution time for the former is approximately linear with increasing sample size, whilst with the latter two the time increase is exponential). TreeTime also has a homoplasy function that can take as input a tree and an alignment, perform ASR and report sites that are homoplasic. TreeTime can report where each mutation at homoplasic SNP sites occur in relation to the tree, but does not attempt to differentiate the various types of homoplasy, nor consider any potential coding consequences of the SNPs. Therefore, if the goal is to find and discard homoplasic SNPs, then HomoplasyFinder or TreeTime are suitable; but if the goal is detailed analysis of mutations potentially contributing to adaptive evolution, further processing of the output is required.

WGS has been widely adopted for the study of bacterial pathogens, and it is frequently applied to datasets numbering in tens to thousands of isolates. WGS is also increasingly used for routine public health surveillance of tuberculosis^17,18^ and foodborne pathogens^19^, generating 10s-100s of thousands of genomes that could potentially be used to interrogate parallel/convergent evolution in natural populations, providing important signals of adaptive evolution via selection for clinically relevant traits such as antibiotic resistance or virulence. There is therefore a need for efficient tools to detect and analyse homoplasic SNPs from large WGS datasets. Such datasets are particularly valuable for selection analysis because they can be highly informative for both phylogenetic tree inference and ASR^20,21,22^ and tend to lead to the recovery of more homoplasic SNPs. However, they also pose substantial computational challenges^22,23^, including generating the phylogenetic tree and mapping the SNPs back to the tree using ASR, as well as post-ASR analysis to extract and classify the homoplasic SNPs. These challenges occur, in part, because both the phylogenetic and ASR inferences require increasingly computationally expensive likelihood calculations for each SNP site (as there are more nodes to resolve over), and because there are typically more SNP sites to test, as the tree size grows. Whilst there have been advances in improving rapid tree inference (*e*.*g*. RAxML-NG^24,25^ and IQTREE^26,27^), ASR (*e*.*g*. TreeTime^15^), and simple detection and filtering of homoplasic sites (HomoplasyFinder^16^ and TreeTime^15^), less attention has been paid to facilitating rapid analysis of convergent evolution from WGS data through detailed analysis of homoplasic SNPs.

Here we present SNPPar, a Python-based program that seeks to speed up the process of homoplasic SNP discovery and analysis in large bacterial WGS data sets (>1,000 isolates and/or >100,000 SNPs). It calls on TreeTime for efficient ASR, and then processes the output to provide a wealth of additional data on mutation events and coding effects to facilitate the detailed assessment of evidence for adaptive evolution in large bacterial SNP data sets.

## 2. Theory and implementation

SNPPar is written in Python 3, utilising the ETE3^28^ package for tree manipulation and TreeTime^15^ or (optionally) FastML^11,12,13^ for ASR. The package is available at https://github.com/d-j-e/SNPPar. Required inputs are a phylogenetic tree (Newick format), annotated reference genome (GenBank format), and details of SNPs for analysis (alignment formats are discussed below). The workflow for SNPPar is outlined in **Figure 2** and summarised below.

**Figure 2.**
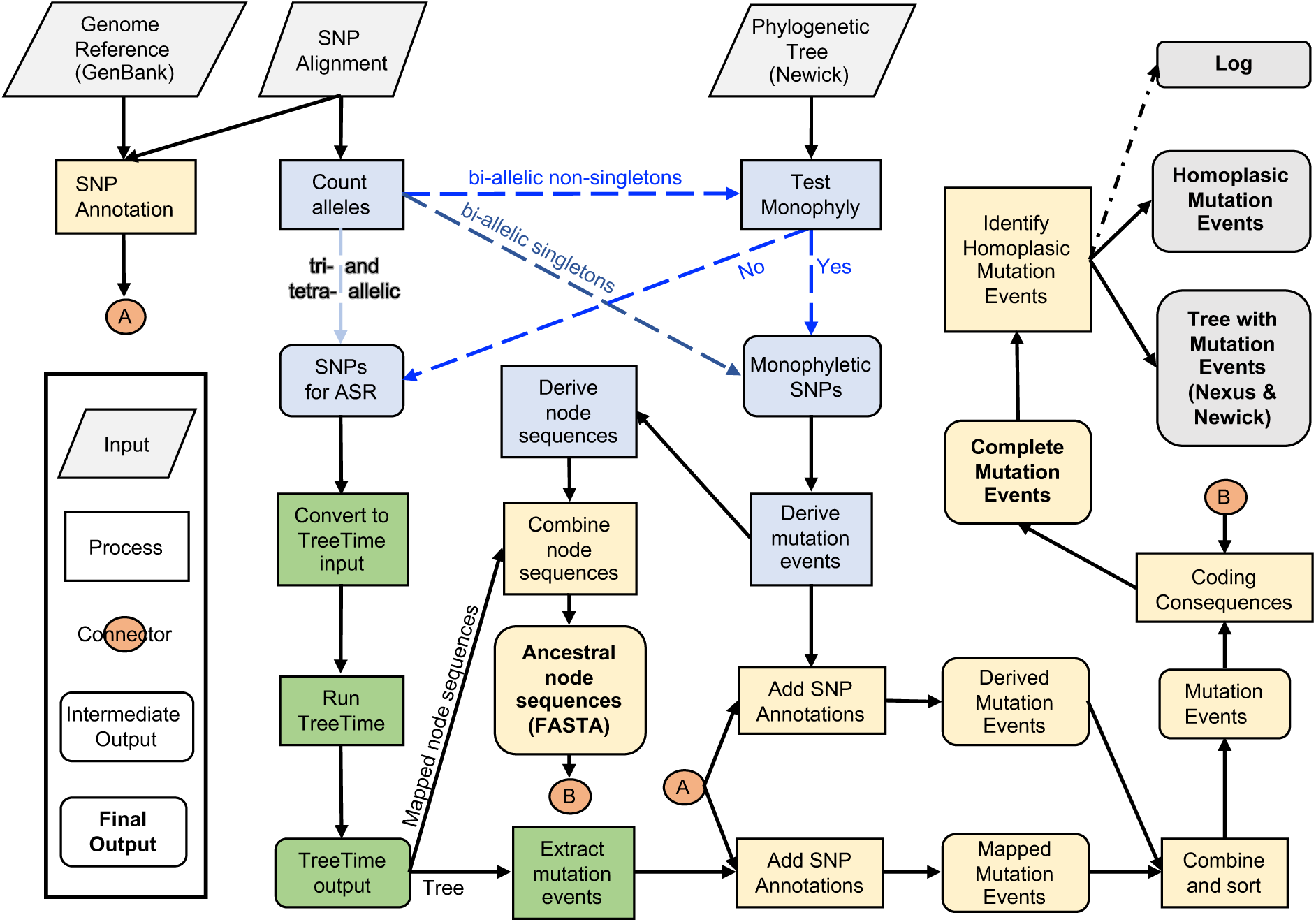
Outline flowchart of SNPPar. The three main components are SNP sorting (blue boxes), ancestral state reconstruction (green boxes), and collation and functional annotation of mutation events (gold boxes). The three different pathways for analysis when sorting SNP sites are indicated by the three sets of dashed arrows (bi-allelic singletons, dark blue; bi-allelic non-singletons, bright blue; tri- and tetra-allelic, pale blue). All outputs are in bold text.

### 2.1 SNP sorting

Firstly, SNPPar sorts SNP sites into different pathways for analysis (see **Figure 2**).

i. Bi-allelic SNPs that are singletons (minor allele present in only one input sequence) are trivially identified as monophyletic and assigned as a single mutation event arising on the relevant terminal branch (see **Figure 3A**).
ii. Bi-allelic SNPs for which the minor allele is present in ≥2 input sequences (*i*.*e*. non-singleton) are assessed against the input tree to determine whether they are monophyletic (and thus assigned as a mutation event arising on the relevant internal branch), or paraphyletic and thus passed to ASR to infer mutation events on the tree (see **Figure 3B-D**). See **Supplementary Information** for details of this step.
iii. Tri-allelic and tetra-allelic SNPs are passed directly to ASR to infer mutation events on the tree.

SNPPar has three different sorting options available, which use all or part of the above pathways. ‘Complex’ sorting employs all of the above steps. ‘Simple’ sorting employs step (i) only and sends all but those SNPs identified as singletons to ASR. ‘Intermediate’ sorting (the default) employs most of the above steps, except any non-singleton biallelic SNP with missing data is sent to ASR without assessing monophyly against the tree, because dealing with missing data can be time-consuming (see below). The resulting SNP set designated for ASR typically includes only a fraction of sites in the input SNP set (∼1% of total SNP sites in the case of complex sorting and ∼40% for simple sorting; intermediate sorting depends on the rate of missingness but falls between the other two). Hence, this sorting stage can considerably reduce processing time compared to conducting ASR on all sites. Initial testing, further confirmed by the validation testing below, led to the conclusion that intermediate sorting for datasets with missing calls was much quicker than the initial implementation, complex sorting, albeit with a small hit to peak memory use (see **Results**). Therefore intermediate sorting was set as the default behaviour for SNPPar, but the other options are still available because (a) simple sorting can be used if users are concerned about the accuracy of the internal monophyly test and prefer to send all non-singleton SNPs to TreeTime, or (b) complex sorting can be used if memory is limited.

**Figure 3.**
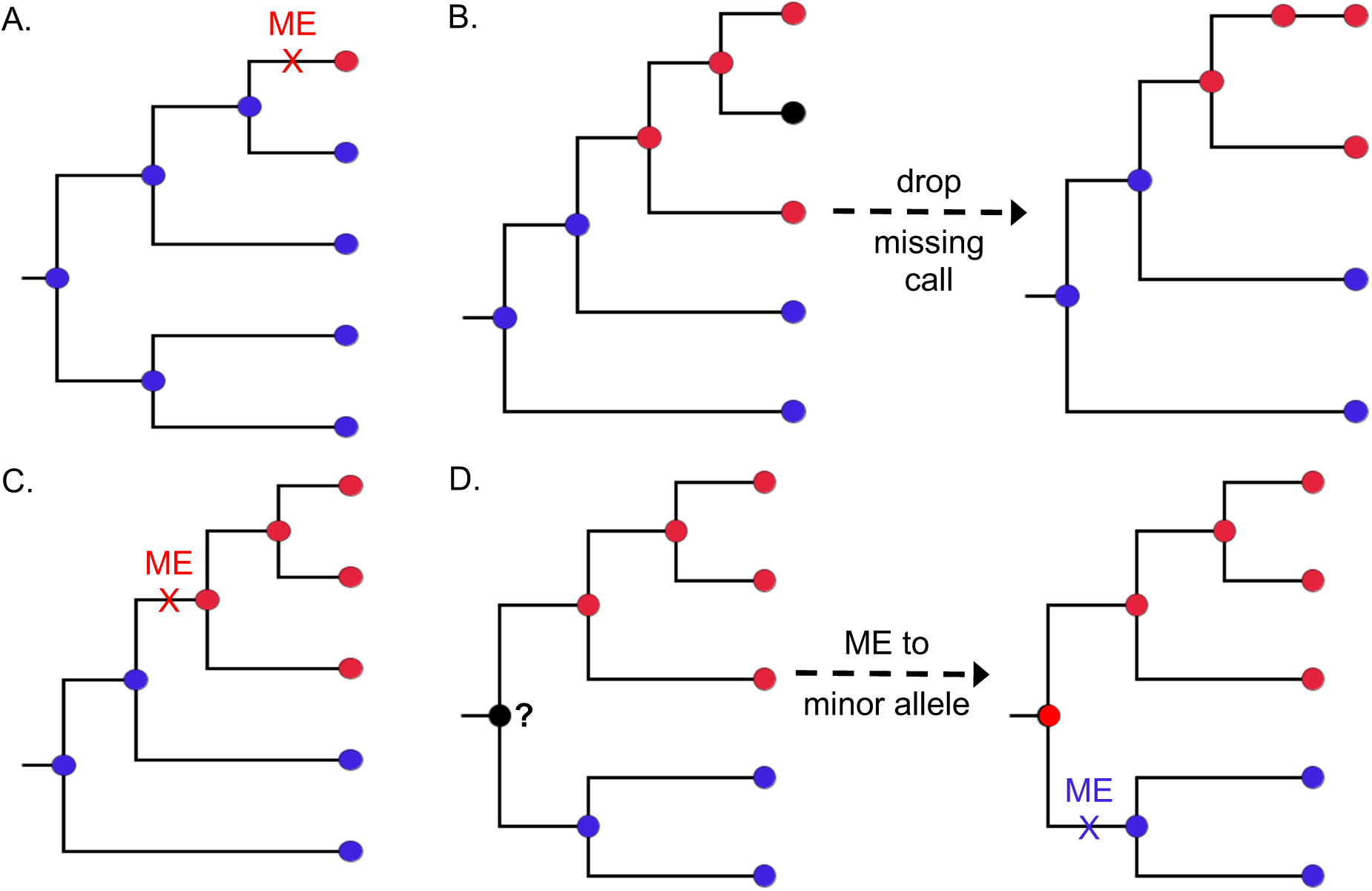
Examples of assigning mutation events for bi-allelic SNP cases. (**A**) singletons. (**B**) Handling missing calls. (**C**) One allele monophyletic. (**D**) Both alleles monophyletic. Each assigned mutation event (ME) is indicated on the branch with a cross, where colour indicates derived base change involved. Note that if neither allele is monophyletic, the site is considered paraphyletic and passed to ASR.

### 2.2 Ancestral state reconstruction (ASR)

ASR is performed for SNP sites identified during sorting, in order to infer mutation events on the tree using a ML approach. By default, SNPPar performs the ASR step by concatenating alleles from the SNP sites selected for analysis into a single FASTA-format alignment, then passing this together with the input tree to TreeTime’s ‘ancestral’ command (note that missing or unknown base-calls can appear in this alignment but are set to ‘N’, as with all maximum-likelihood methods). TreeTime’s default behaviour is to ignore sites with any missing base-calls, hence SNPPar uses the TreeTime option to enforce reporting of all sites. Two output files are captured for processing by SNPPar: the sequences reconstructed for each internal node in the tree (‘node sequences’), and a copy of the input tree with the inferred mutation events annotated on each branch (‘event tree’), from which SNPPar extracts a list of all mutation events per SNP site. Optionally, FastML can be selected as an alternative ML algorithm for ASR, however this is much slower, and we found no difference in accuracy (see **Results**).

### 2.3 Collation and functional annotation of mutation events

All mutation events, whether inferred via internal sorting or ASR, are collated into a master event list that records the SNP coordinate in the reference genome, the precise base substitution (ancestral to derived base), and branch for each mutation event. These events are then annotated with their functional effects, with reference to the protein-coding gene (CDS) features annotated in the input reference genome and taking into account the inferred codon at each node not just the SNP site (see **Supplementary Information** for details). The final step in SNPPar is to identify, count and report unique homoplasic SNPs found in the full mutation event list. SNPPar can also identify the type of homoplasy for each homoplasic SNP (*i*.*e*. parallel, convergent and/or revertant mutation events), either in the main analysis run or as a post-processing step applied to a previously generated mutation event list and tree.

### 2.4 Input and output files and formats

The annotated reference genome must be in GenBank format. SNPPar assumes the reference is circular (for the purpose of reporting relative position of intergenic SNPs), and ignores any CDS that are split across the origin. The input tree must be rooted, bifurcating and in Newick format, with branch lengths expressed in substitutions per site. SNPs can be input in two ways: (i) a multi-FASTA concatenated alignment of SNP alleles (FASTA headers being sample names that match exactly the tip names in the input tree) plus a list indicating the coordinates of SNPs relative to the reference genome (one SNP coordinate per line, in the order in which SNP alleles appear in the alignment); or (ii) a comma-separated SNP allele table, with SNPs in rows (column 1 indicating the SNP coordinates, relative to the reference genome) and samples in columns (column headings being sample names that match exactly the tip names of the input tree).

Primary outputs are an annotated list of all mutation events and their coding effects, an annotated list of just those mutation events found at homoplasic SNP positions, and two tree files in which nodes are labelled with identifiers and mutation event counts (total SNPs and homoplasic SNPs). One tree file is in NEXUS format, compatible with visualisation in FigTree (http://tree.bio.ed.ac.uk/software/figtree/) or iTOL^29^; the other contains the same data but in extended Newick format, suitable for use with the R package ggtree^30^. Internal node sequences are also output as multi-FASTA alignments. All program events are recorded to a time-stamped log file, including any relevant warnings (*e*.*g*. split gene in GenBank reference) or critical errors (*e*.*g*. providing FASTA alignment, but not the SNP position list).

## 3. Methods for validation and performance testing

All development and testing was done on a 2017 Apple MacBook Air (1.8 GHz Intel Core i5, 8 GB of RAM) using Python v3.7.2, ETE3 v3.1.1 and TreeTime v0.6.3. Results using FastML v3.11 for ASR on simulated data are included for comparison purposes. The full protocol, scripts and data used to carry out the validation and performance testing are available on GitHub (https://github.com/d-j-e/SNPPar_test).

### 3.1 Validation with simulated data

In order to test the accuracy of SNPPar when performing homoplasic SNP detection, we simulated 120 sets of sequences based on substitution model parameters and phylogenetic trees estimated from real *Mycobacterium tuberculosis* (*Mtb*) SNP alignment data^31^. Note this SNP data had been filtered previously to mask repetitive regions of the genome.^31^ The data was divided into three subpopulations representing distinct epidemiological scenarios: (1) single lineage, single location (821 *Mtb* lineage 2 isolates from Ho Chi Minh City; HCMC L2); (2) single lineage, globally distributed (940 *Mtb* lineage 2 isolates; Global L2); and (3) species-wide, globally distributed (2965 isolates from lineages 1, 2 and 4; Global L124). Empirical parameters were extracted from 10 randomly sampled subsets of various sizes from each population (10%, 20%, 50% and 100% of isolates from each of HCMC L2 and Global L2; n=100, 500, 1,000 and 2,000 isolates from Global L124). For each subset, (i) the number of SNPs was recorded and used later to correct the number of SNPs generated by simulation; and (ii) a reference tree and GTR model parameter estimates (alpha, base mutation frequencies and nucleotide frequencies) were obtained by selecting the tree with the highest likelihood from a random sample of five trees estimated from the SNP alignment using RAxML v8.2.12 using a GTR+G model and ascertainment bias correction. SeqGen v1.3.4^32^ was used then used to simulate sequence evolution along these trees (see details in **Supplementary Information** and the SNPPar_test GitHub repository).

SNPPar was run on each simulated data set using either simple and intermediate sorting, with homoplasic SNP classification turned on. (Note as there are no missing SNP alleles in the simulated data, complex sorting would be identical to intermediate sorting.) The list of homoplasic SNPs output by SNPPar was compared to the list of those expected based on the simulation. The starting tree was used as input rather than inferring new trees from each simulated alignment, because this would test the accuracy of the tree-building stage as much as the ability of SNPPar to find homoplasic SNPs. Any SNP expected to be called as homoplasic that was reported as such by SNPPar was scored as a true-positive; any other homoplasic SNP reported was considered a false-positive; any expected homoplasic SNP not reported by SNPPar was considered a false-negative. For homoplasic SNP calls, we also considered whether the type was classified correctly (*i*.*e*. parallel, revertant or convergent). As only one case of convergence was expected (and was always detected), only parallel and revertant type calls were considered.

SNPPar’s total run time and maximum memory usage was recorded for each replicate run. The results were analysed in R v3.5.1 using RStudio v1.1.383, with independent ANOVA tests to examine the effects of isolate count, SNP count or total alignment length (*i*.*e*. total isolate count multiplied by SNP count) on total run time and memory.

### 3.2 Performance validation with real data

Testing of the performance of SNPPar in real-world analysis scenarios was demonstrated using two previously published bacterial WGS data sets in which convergent SNPs have been examined. One was from a 2015-16 *Elizabethkingia anophelis* outbreak in Wisconsin, USA^33^ (68 isolates, 369 SNPs), the other from a retrospective sequencing study of in-host evolution and transmission of *Burkholderia dolosa* isolates amongst cystic fibrosis patients receiving treatment at a hospital in Boston, USA, in the 1990s^34^ (114 isolates, 511 SNPs). As *B. dolosa* have three chromosomes, each reference chromosome and its associated SNPs were analysed separately using SNPPar (299, 153, and 59 SNPs on chromosomes 1, 2 and 3, respectively). The published tree for *B. dolosa* had branch lengths in units of substitutions (i.e. SNP counts), hence these were scaled to units of substitutions per site by dividing each by the chromosome sequence length prior to use as input for SNPPar.

The performance of SNPPar in real-world analysis scenarios was further demonstrated using the largest subsampled empirical data set, Global L124 with 2000 isolates. To this group of samples, we added an outgroup (a single Lineage 5 isolate) and extracted the SNPs as previously above. We then constructed the phylogeny using IQTREE 2^26,27^ using a transversion model for the mutation matrix, as indicated by IQTREE, and rooted it using the outgroup. The resulting tree was used in SNPPar along with the SNP dataset to obtain the mutation events across all three lineages (*i*.*e*. excluding the outgroup). Before analysis with SNPPar, we identified genes that overlap with masked repetitive regions, removed from the SNP alignment all SNPs located in these genes, and excluded these genes from further downstream analysis.

## 4. Results

### 4.1 Validation of parallel SNP detection using simulated data

We assessed accuracy of SNPPar using *Mtb* data simulated for three different population structures (single lineage, single city; single lineage, global; species-wide, global), subsampled at four different sizes, each with 10 replicates (total 120 simulations; see details in **Methods**). The two key outcomes assessed were the correct detection of homoplasic SNPs, and the correct identification of the type of homoplasy for each reported homoplasic SNP. The effects of sample characteristics (population structure, sample size and SNP count) on the two key outcomes was also explored.

#### 4.1.1 Accuracy

Accuracy results (using the default option of intermediate sorting) are summarised in **Figures 4-5**. Note the results produced by SNPPar are deterministic; running the same inputs more than once produces exactly the same outputs. This extends to the choice of sorting; amending how SNPPar sorts the SNPs has no effect on accuracy when there is no missing data (as with the simulated datasets), and very minor effect if there is missing data.

**Figure 4.**
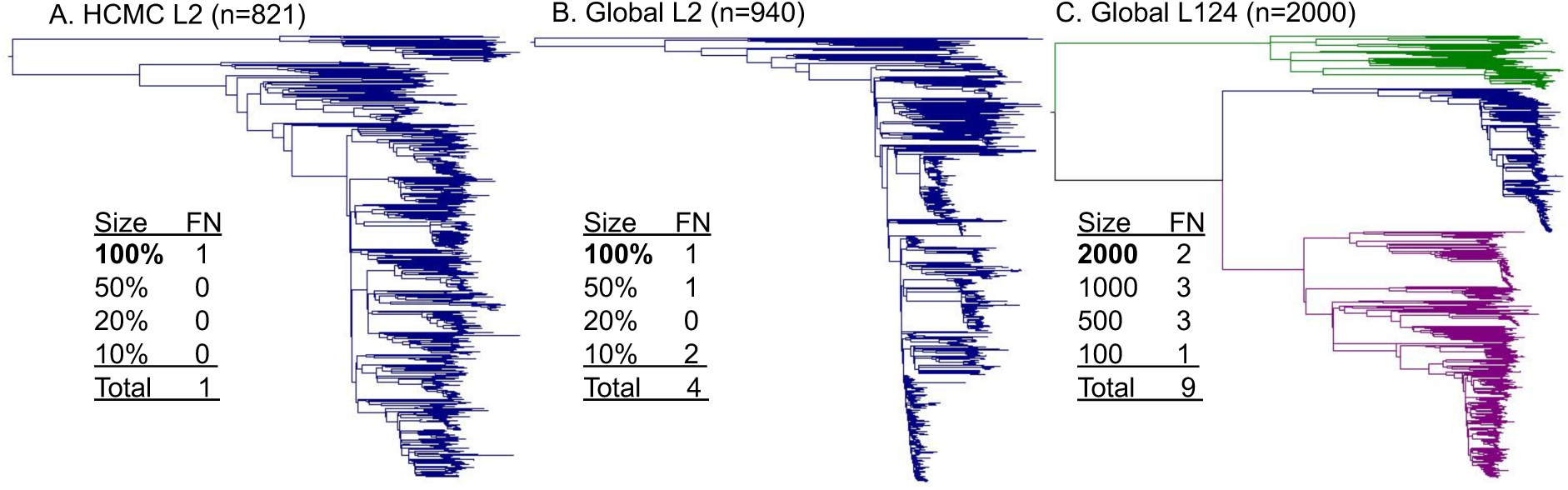
Example of the largest trees from the three populations used for simulating *M. tuberculosis* data sets, A) HCMC L2; B) Global L2; and, C) Global L124. The tree in (**C**) consists of three lineages of *M. tuberculosis*; lineage 1 (green), lineage 2 (blue), and lineage 4 (grey), whilst (**A**) and (**B**) consist of only lineage 2 isolates. Inset tables show the number of false-negative (FN) homoplasy calls found in each set of 10 replicate simulations conducted for each sample size for each population (bold indicates tree size illustrated). FN calls are defined as homoplasies present in the simulated datasets that were not detected by SNPPar. Sample sizes for the HCMC L2 and Global L2 populations are random samples of genomes representing a set percentage subsample of the total sample size; for Global L124, a fixed number of genomes was sampled. Note there were no false-positive homoplasy calls across all simulated datasets.

SNPPar generated no false-positive homoplasy calls across the 120 replicate tests. Most replicate runs produced either zero (n=107 runs) or one (12 runs) false negatives; the single exception was one run on a dataset of the smallest sample size from the Global L2 population, in which two false negative calls were made. As a general trend, the more complex the population, the more likely a homoplasy would be missed (**Figure 4**). Notably, in all cases it was more likely to encounter no false negatives at all, *i*.*e*. all homoplasic SNPs were detected. Three types of false-negative homoplasy calls were observed; all involved SNPPar failing to identify a homoplasy in a scenario for which the available data supported a simpler single-mutation explanation (*e*.*g*. parallel mutation events on sister branches, which were called as a single mutation on the branch leading to the parent node; **see Figure S1A-C**). Note that such scenarios are also not possible to detect using alternative approaches (such as ASR on all sites, or HomoplasyFinder’s consistency index).

The true-positive homoplasic SNPs were assessed to see whether the type (parallel, convergent or revertant) was correctly reported by SNPPar. Overall, most types were correctly identified, and accuracy improved with sample size (see **Figure S2A**). In the smallest sample size (n=100), a mean of 13.7 homoplasic SNPs were called, of which mean 8.4% were classified incorrectly; in the largest samples (n=2000), a mean 425.5 homoplasic SNPs were called of which mean 0.84% were classified incorrectly. The most common error was a reversion being incorrectly classified as parallel event), followed by a parallel event being incorrectly classified as a reversion (see **Figure S2B**); both cases were caused by the necessity to make arbitrary ancestral allele calls at the root node, which is unresolvable with any method.

Using FastML for the ASR step yielded no improvement in accuracy over TreeTime, producing the same false-negative calls and no false-positive calls. FastML yielded some differences in homoplasy type call errors, with a higher overall error rate (∼5% more incorrect type calls with the smallest data set and ∼0.7% more errors with the largest, see **Figure S3A**), but this was due solely to differences in arbitrary assignments at the root node (see **Figure S3B**).

#### 4.1.2 Performance

Resource usage (total run time and peak memory) for SNPPar analyses with empirical and simulated data are shown in **Figure 5**. For the simulated datasets sorting options included simple and intermediate (as there is no missing data), whilst all three options were tested with the empirically subsampled (real) datasets.

**Figure 5.**
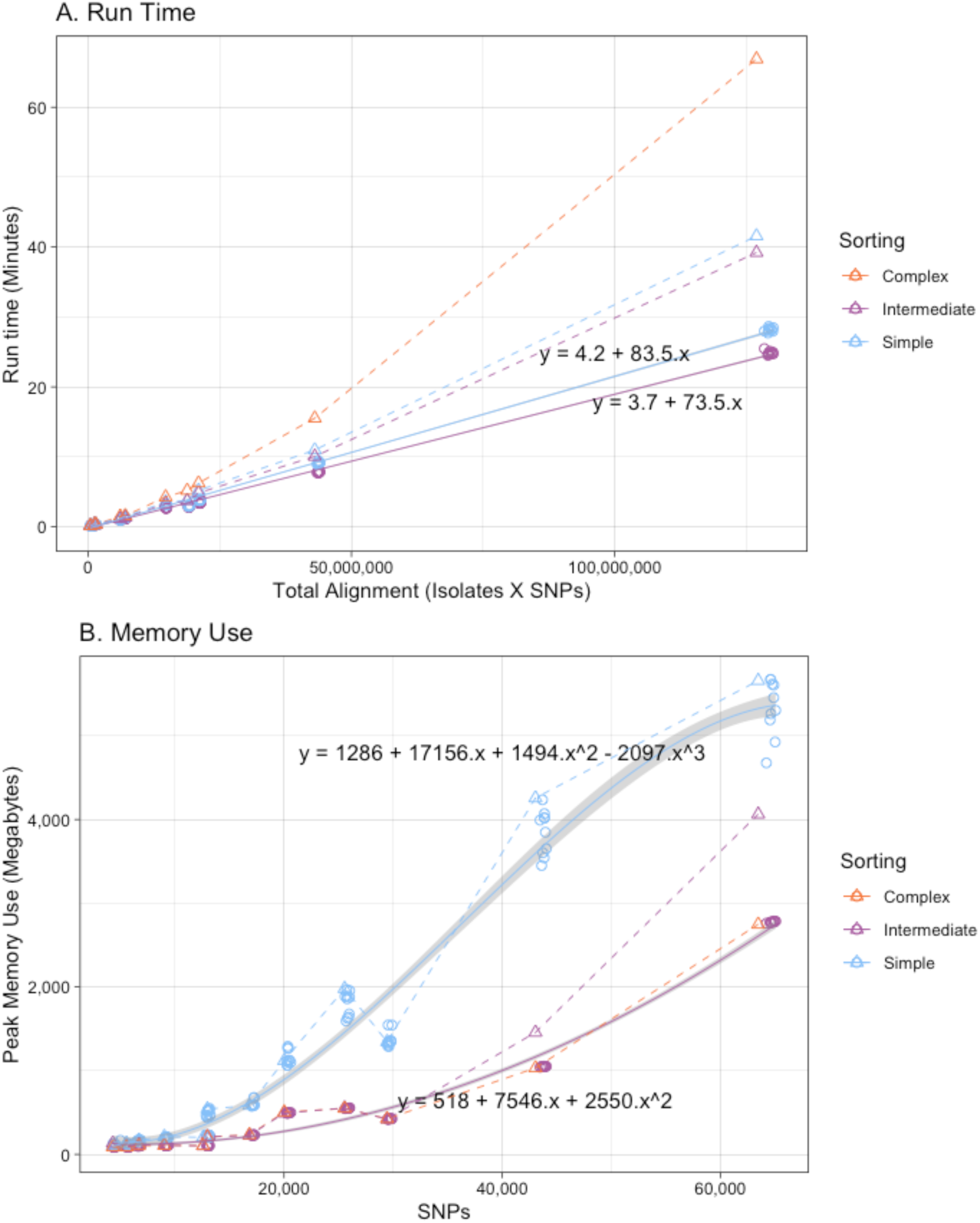
SNPPar performance during analysis of subsampled empirical and simulated datasets using different sorting options. (**A**) Total run time vs total alignment length. (**B**) Peak memory use vs SNP alignment length. Circles denote simulated datasets, triangles empirical datasets; colours indicate sorting options as per inset legend. The simulated datasets have ten replicates of each sample size, and a line of best fit through these is indicated (solid line, grey shading indicates 90% fit interval). Note complex sorting was not used for the simulated datasets as there are no missing calls, and complex and intermediate sorting are identical in this case. The real datasets have only a single observation per sorting algorithm, which are joined with simple dashed lines.

Variance in total run time was most strongly associated with total alignment size of the input dataset, whilst the peak memory use was most strongly associated with number of SNPs (see **Supplementary Information**). Whilst the choice of sorting algorithm for the simulated datasets had only a minor impact on total run time (intermediate being slightly quicker than simple sorting, solid lines in **Figure 5A;** ∼11% quicker for intermediate sorting with the largest data set, Global L124 with a sample size of 2000), there was a major difference in the peak memory use as more SNPs were sent to ASR with simple sorting (solid lines in **Figure 5B**; ∼49% more memory used for intermediate sorting with the largest data set). Memory use followed the same pattern for the empirical data (which includes missing alleles and many more homoplasic SNPs), with simple sorting (most SNPs sent to ASR) using the most memory and complex sorting (fewest SNPs sent to ASR) the least (dashed lines in **Figure 5B**). However, this comes at the expense of runtime; empirical datasets took longer to analyse than similar-sized simulated datasets, and complex sorting took ∼50% longer than simple or intermediate sorting (dashed lines in **Figure 5A**). Hence, whilst the default setting of intermediate sorting was consistently the fastest (∼40 minutes on the largest empirical dataset), this increase does come at a cost to peak memory use compared to the slower complex sorting (∼4 GB vs ∼2.5 GB on the largest empirical dataset), which may become problematic with very large datasets.

Comparing the performance of SNPPar to TreeTime with regards to the ASR step, the whole alignment of a simulated data set of Global L124 with 2,000 isolates took ∼22 minutes for TreeTime to analyse all ∼64000 SNPs using ∼24 GB peak memory; SNPPar using the default intermediate sorting sent only ∼800 SNPs (1.25% of the total) to TreeTime for ASR, which took ∼23 seconds with a peak memory use of ∼0.4 GB of RAM.

Using FastML rather than TreeTime for ASR in SNPPar saw a minor improvement on memory usage but increased the run time markedly, especially for larger data sets (see **Figures S4A and B**). For example, the largest simulated data set (Global L124 with 2,000 isolates) took on average ∼25 minutes and used ∼2.7 GB of memory with TreeTime (using intermediate sorting) and ∼3 hours and ∼1.6 GB with FastML. FastML by itself, given the same whole alignment of Global L124 simulated data and computing resources as TreeTime above, failed to complete the analysis after seven days. SNPPar using FastML also failed (*i*.*e*. exited with a fatal error) during the ASR step for two of the largest real SNP alignments (Global L124 with 2,000 isolates, 1254 SNPs sent to ASR; and HCMC L2 with 821 isolates, 395 SNPs sent to ASR) which completed in ∼25 and ∼3 minutes, respectively, using SNPPar with TreeTime and intermediate sorting).

### 4.2 Demonstration of SNPPar use cases with real data

Two data sets from previously published papers reporting homoplasy analysis in *E. anophelis*^33^ and *B. dolosa*^34^, along with the largest empirically sampled *Mtb* dataset used for simulation (Global L124 with a sample size of 2,000 plus outgroup), were analysed with SNPPar using default parameters (details in **Methods**). The data sets, commands and outputs are available on GitHub (https://github.com/d-j-e/SNPPar_test).

*E. anophelis* is an opportunistic pathogen that rarely causes outbreaks, and the genome data from an unusually large outbreak in the US provide a rare opportunity to explore positive selection in this organism.^33^ SNPPar analysis of the alignment of 369 SNP sites from 68 *E. anopheles* outbreak genomes took <3 seconds, and identified the same homoplasic SNP that was reported in the original study (which used FastML)^33^ (see **Figure 6**). The SNP (G to T at position 2,169,660 of the reference sequence) was assigned to two different terminal branches, indicating it had arisen via parallel substitutions in two different outbreak strains (CSID3000521203 and CSID3015183676). SNPPar correctly annotated this as a parallel SNP affecting the first position of codon 469 in the *wzc* capsular export gene (A2T74_09840), resulting in a nonsense mutation (converting the glutamate codon ‘GAA’ to stop codon ‘TAA’) and thus truncating the protein as previously reported.^35^ Because SNPPar annotates all mutation events, the output can also trivially be used to extract gene-level information about selection, not just homoplasic SNPs. For example, the original study^33^ reported that 27 genes had ≥2 independent protein-altering mutations (*i*.*e*. nonsynonymous or nonsense mutation events) in the outbreak population, including three with ≥5 protein-altering mutations (a hypothetical protein potentially involved in starch utilization, the capsular export gene *wzc*, and another sugar transporter similar to *wza*). SNPPar identified the same numbers of mutation events in these 27 genes. Five protein-altering SNPs (involving six independent mutation events) were correctly found in *wzc*, including the two for the parallel nonsense SNP at codon 469 (see **Figure 6**).

**Figure 6.**
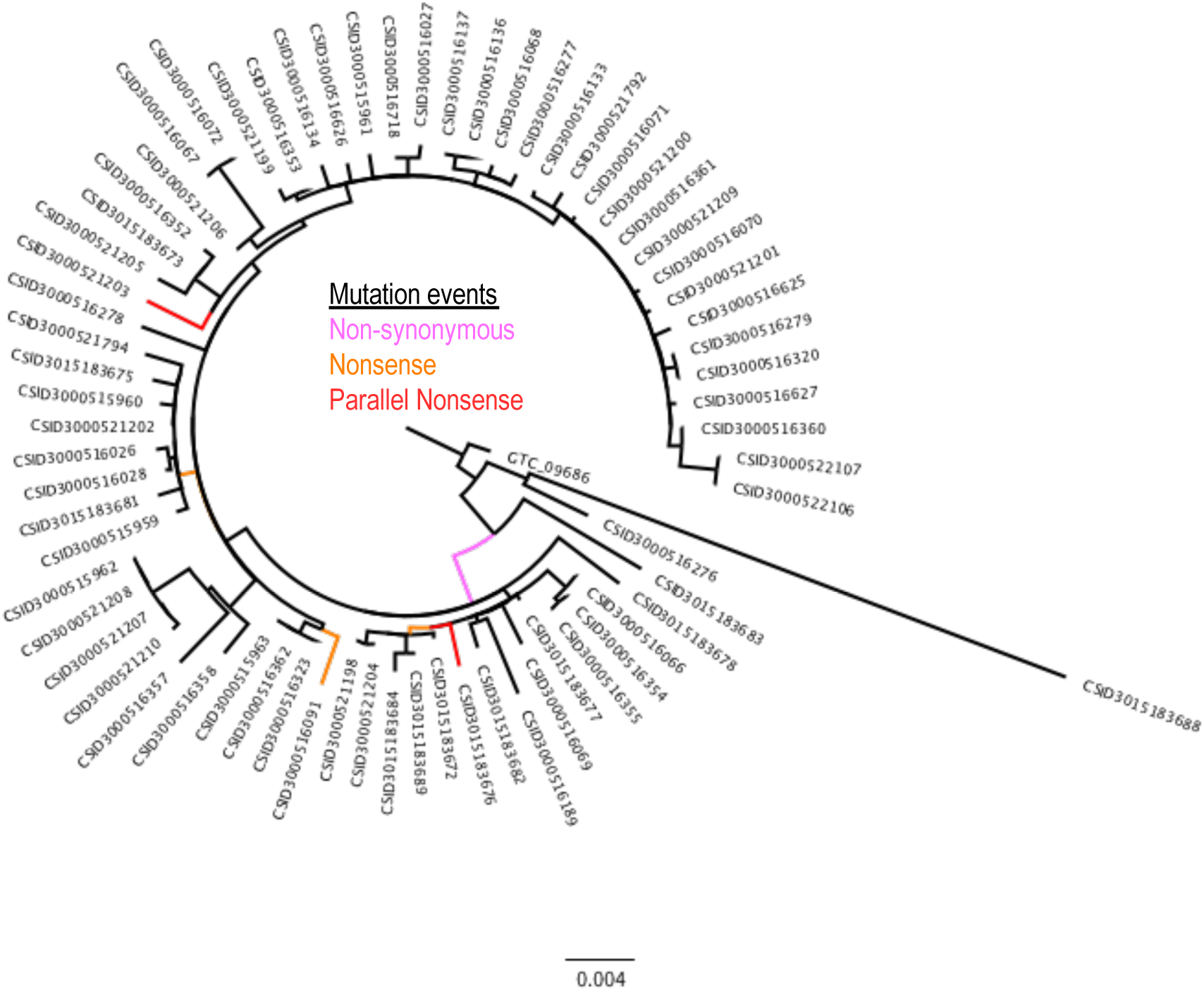
Real data example: 2016-17 *Elizabethkingia* outbreak maximum likelihood phylogeny^33^ showing location (branch) of the six protein-altering mutation events of the *wzc* capsular export gene. The three types of mutation event shown are nonsynonymous (one event, pink), nonsense (three events, orange) and parallel nonsense (two events, red). The tree includes an outgroup isolate (GTC_09686).

WGS from serial *B. dolosa* isolates from chronically infected cystic fibrosis patients provides an opportunity to investigate in-host evolution of the pathogen^34^. Under these conditions the bacterial population is constrained by isolation and strong selective pressure from the host, hence substitutions are a dominant mechanism of adaptation and parallel mutations can provide a strong signal of positive selection. *B. dolosa* has three chromosomes and SNPPar was run on separate alignments for each chromosome (using a single consensus tree of 114 isolates inferred from the concatenated alignment of 511 SNPs), which took a total of ∼11 seconds. Twenty SNP sites with >1 mutation event were previously reported in this data set^34^; SNPPar correctly identified all twenty (see **Table 1**), despite the alignment having high levels of missing alleles for some SNPs (up to 106/114 alleles missing at a given SNP site). The SNPPar output includes the inferred type of homoplasy for each position, along with the precise mutation event/s involved and annotation of the coding effects (see **Table 1**). Fourteen intragenic parallel SNPs were detected, notably all were nonsynonymous. Three genes harboured multiple nonsynonymous parallel SNPs, consistent with strong positive selection for modified protein function (highlighted in **Table 1**). All three of these genes were associated with antibiotic resistance. Parallel and divergent SNPs were detected in codons 83 and 87 of *gyrA* (BDAG_02180; see **Figure 7A**), these were linked to fluoroquinolone resistance in the original study^34^. Two distinct parallel SNPs were identified in codon 172 of beta-lactamase *bla1* (BDAG_03694), consistent with the original study in which this gene was noted as under positive selection^34^. The third gene was the ribosomal 50S L4 protein-encoding gene *rpl4* (BDAG_02781), which is involved in both pathogenicity and macrolide resistance^36^. SNPPar identified parallel SNPs in the first and second base positions of codon 70 of *rpl4*, consistent with the original report which identified it as under positive selection, with 4 independent mutations affecting codon 70^36^. As shown in **Figure 7B**, SNPs at these two positions in *rpl4* arose in parallel a number of times (4 and 8 respectively), but never in the same isolate. Overall, SNPPar identified five parallel nonsynonymous SNPs that arose three or more times independently in the population, which we take to indicate a strong signal of positive selection: one *gyrA*-83 SNP, one *bla1*-172 SNP, the two *rpl4* SNPs, and a SNP in codon 275 of glycosyltransferase gene *wbaD* (BDAG_02317). Notably an additional divergent SNP also occurred at the same site in *wbaD* codon 275 (see **Table 1**). As noted in the original study, the ancestor of this *B. dolosa* cluster harbours a premature stop codon at this position, and the SNPs at this site restore the truncated form of the protein to the full-length protein, restoring proper function of the gene and generating O-antigen repeats. SNPPar results are consistent with the analysis conducted in the original study (see **Figure 7C**), which reported that restoration of the protein via SNPs at this site occurred independently in nine subjects^34^.

**Table 1.** Summary of SNPPar results for *B. dolosa* loci in which homoplasic or divergent SNPs were identified^34^. Bolded genes are those with parallel SNPs that occur ≥3 times independently and are discussed in the text. The type of change (coding effect) of SNPs can be synonymous (S) or nonsynonymous (NS). Amino acid changes are indicated in the format [ancestral amino acid, codon, derived amino acid], *e*.*g*. P537L indicates that the amino acid at codon 53 has changed from proline (P) to leucine (L); * indicates a stop codon.

| Locus Tag | Chromosome<br>::Position | Mutation<br>Events | Type of<br>Mutation Event(s) | Base<br>Substitutions | Type of<br>Change | Amino Acid<br>Change |
| --- | --- | --- | --- | --- | --- | --- |
| BDAG_00179 | 1::232898 | 2 | 1 reversion | C to T, then<br>T to C | NS | P537L then L537P |
| BDAG_01022 | 1::1267993 | 2 | 1 parallel | A to C | NS | K30T |
| intergenic | 1::1431059 | 2 | 2 divergent | C to G<br>C to T | -<br>- | -<br>- |
| **BDAG_02180**<br>**(gyrA)** | 1::2694721 | 4 | 2 parallel | G to A | NS | D87N |
|  |  |  | 1 divergent | C to G<br>G to T | NS<br>NS | H87D<br>D87F |
|  | 1::2694732 | 6 | 4 parallel | C to T | NS | T83M |
|  |  |  | 1 divergent | C to A | NS | T83K |
| **BDAG_02317**<br>**(wbaD)** | 1::2849264 | 10 | 8 parallel | T to G | NS | *275E |
|  |  |  | 1 divergent | T to C | NS | *275Q |
| BDAG_02593 | 1::3170526 | 2 | 2 divergent | T to C and T to G | both S | - |
| intergenic | 1::3380836 | 2 | 2 divergent | G to T and G to C | - | - |
| **BDAG_02781**<br>**(rpl4)** | 1::3390724 | 5 | 3 parallel | G to C | NS | G70A |
|  |  |  | 1 divergent | G to A | NS | G70D |
|  | 1::3390725 | 9 | 7 parallel | G to C | NS | G70R |
|  |  |  | 1 divergent | G to A | NS | G70S |
| BDAG_03003 | 2::261481 | 2 | 1 parallel | A to C | NS | K1525T |
| BDAG_03043 | 2::330177 | 2 | 1 parallel | T to G | NS | S28R |
| BDAG_03095 | 2::397597 | 2 | 1 parallel | T to G | NS | L35V |
| intergenic | 2::701457 | 2 | 1 parallel | A to C | - | - |
| intergenic | 2::819544 | 2 | 1 parallel? | T to G, or<br>G to T, then<br>T to G | - | - |
| **BDAG_03694**<br>**(bla1)** | 2::1231197 | 6 | 3 parallel | C to A | NS | R172S |
|  |  |  | 1 parallel | C to G | NS | R172G |
| intergenic | 2::1616780 | 3 | 2 parallel | G to A | - | - |
| BDAG_04180 | 2::1838491 | 3 | 2 parallel | C to T | NS | A103V |
| BDAG_04506 | 3::70968 | 2 | 1 parallel | C to G | NS | H209D |
| intergenic | 3::72093 | 2 | 1 parallel | C to A | - | - |

**Figure 7.**
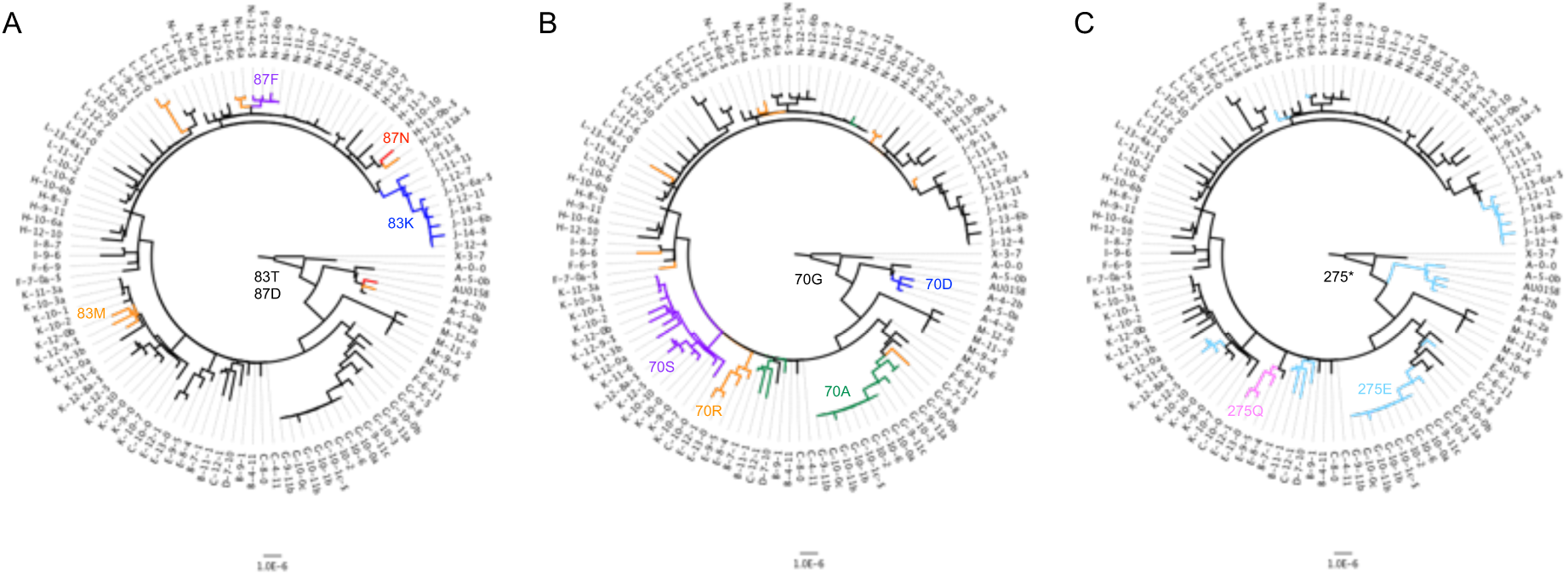
Real data example: *B dolosa* phylogeny^34^ showing evolution of codons 83 and 87 in *gyrA* (A), codon 70 in *rpl4* (B), and codon 275 in *wbaD* (C). Scale bar given in substitutions/site. The tree includes an outgroup (X-3-7) from prior to the transmission chain. Labels for isolates from the same patient start with the same capital letter. Encoding of the codon(s) at any position in the tree is indicated by branch color with the ancestral call indicated with black. In **A**, the ancestral codon 83 of gyrA that encodes to threonine (T) is changed to methionine (M, orange), phenylalanine (F, purple) or lysine (K, blue). Codon 87 in the same gene changes from aspartic acid (D) to asparagine (N, red). In **B**, the ancestral codon 70 of *rpl4* that encodes to glycine (G) changes to serine (S, purple), arginine (R, orange), alanine (A, green) or aspartic acid (D, blue). In **C**, the ancestral (truncating) stop codon of *wbaD* (position 275) is changed to either glutamic acid (E, light blue) or glutamine (Q, pink).

For the *Mtb* dataset (total 63,065 SNPs in 2000 genomes), SNPPar took 51 minutes (using 2.6 GB) and identified a total of 2879 homoplasic mutation events affecting 461 genes. We used the output to calculate the rate of homoplasic protein-altering mutations per gene, normalised to total number of synonymous mutation events per gene, as a measure of adaptive selection (see **Figure 8A)**. The top ten scoring genes (see **Table 2**) include five known targets of positive selection associated with antibiotic resistance^35,36^ (*rpsL*^37^, *katG*^38^, *pncA*^39^, *rpoB*^40^ and *embB*^41^). The other five encode hypothetical proteins, though one of these (*sseC2*) has been implicated in down-regulation of genes involved in latency in *Mtb*.^42^ The five antibiotic resistance-associated genes each harboured homoplasies at ≥3 nucleotide positions (from three in *katG* and *rpsL*, across two and three codons respectively, to 18 in *pncA*, across 18 different codons; see **Figure 8B**). In *katG*, which is known for its isoniazid-resistance conferring mutations^38^, most of the homoplasic mutation events (90/96) occurred in codon 315 (see **Figure 9A**). The majority of these (80/90) resulted in the same amino acid substitution (S315T), though there were also three (parallel) reversions at the same codon position (i.e. T315S, largest affected clade shown in **Figure 9B**). Note that these exploratory analyses are simple to conduct using the tabular and tree output of SNPPar.

**Table 2.** Top ten *M. tuberculosis* genes in terms of ratio of protein-altering homoplasic mutation events vs all synonymous mutation events. Bold gene names indicate genes previously reported by Fahrat et al.^35^ and Coll et al.^36^ to be targets of convergent positive selection. ME, mutation event count; S, synonymous SNP count; PA, protein-altering mutation count; sites, number of nucleotide sites affected by ≥1 homoplasic SNP.

| Gene_Tag | ME | S | PA | PA/(S+1) | Sites | Gene | Product | Function (Reference) |
| --- | --- | --- | --- | --- | --- | --- | --- | --- |
| Rv0682 | 76 | 3 | 73 | 14.6 | 3 | **rpsL** | 30S ribosomal protein S12 | streptomycin resistance (37) |
| Rv1908c | 96 | 0 | 96 | 13.7 | 3 | **katG** | catalase-peroxidase | isoniazid resistance (38) |
| Rv2043c | 45 | 0 | 45 | 11.3 | 18 | **pncA** | pyrazinamidase/nicotinamidase | pyrazinamide resistance (39) |
| Rv0336 | 21 | 0 | 21 | 10.5 | 2 | - | hypothetical protein | - |
| Rv0667 | 140 | 0 | 140 | 9.3 | 8 | **rpoB** | DNA-directed RNA polymerase subunit beta | rifampin resistance (40) |
| Rv2828A | 17 | 0 | 17 | 8.5 | 1 | - | hypothetical protein | - |
| Rv1944c | 16 | 0 | 16 | 5.3 | 2 | - | hypothetical protein | - |
| Rv0814c | 5 | 0 | 5 | 5.0 | 1 | **sseC2** | hypothetical protein (sulphur metabolism) | latency (42) |
| Rv3795 | 124 | 0 | 124 | 4.6 | 10 | **embB** | arabinosyltransferase B | ethambutol resistance (41) |
| Rv0750 | 52 | 8 | 44 | 4.4 | 8 | - | hypothetical protein | - |

**Figure 8.**
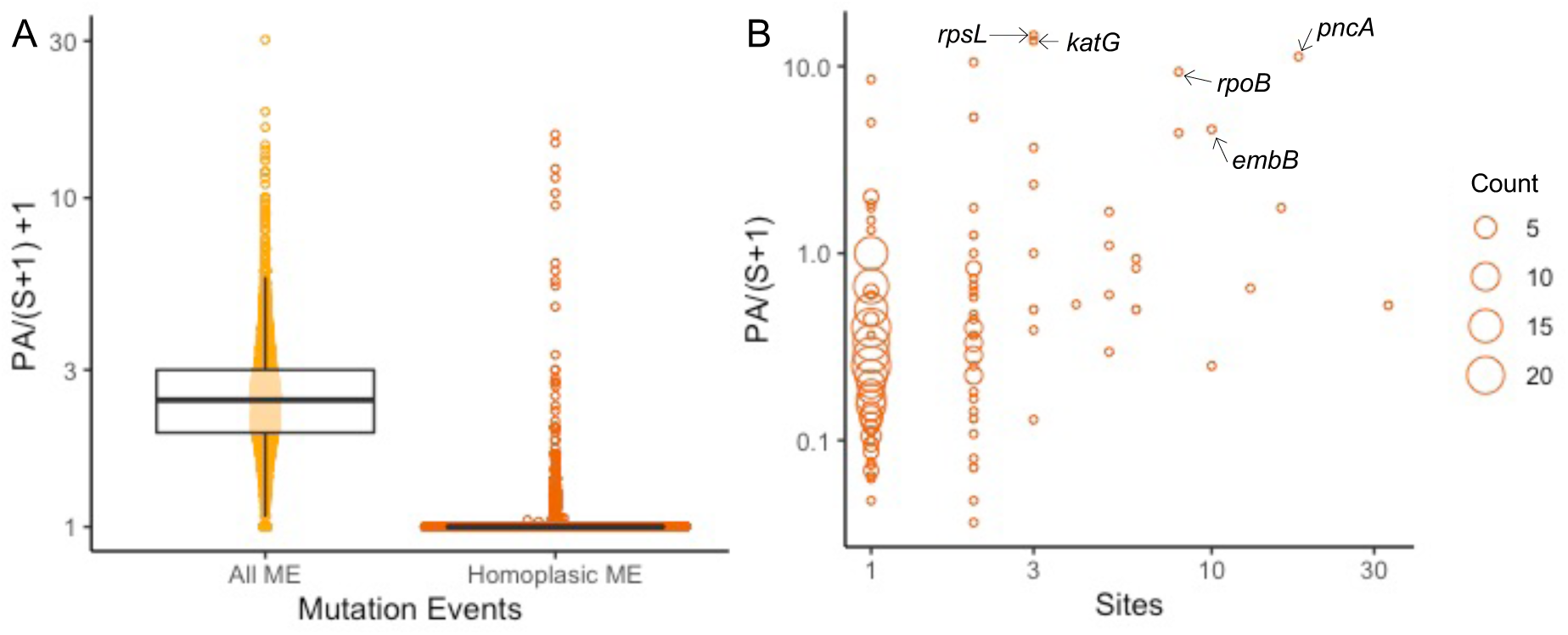
Real data example: Gene-level summary of homoplasies detected in *Mycobacterium tuberculosis*. (**A**) Distributions for ratio of protein-altering (PA) events per gene normalised to synonymous SNP count per gene, for all (yellow) and homoplasic (orange) mutation events. (**B**) Number of nucleotide sites affected by homoplasies (x-axis) vs PA ratio (y-axis), each point represents one or more genes with the same (x,y) values, circle size indicates the number of genes (as per inset legend). Resistance genes discussed in the text are labelled. Note that this analysis includes 2000 strains randomly sampled from the Global L124 set^31^.

**Figure 9.**
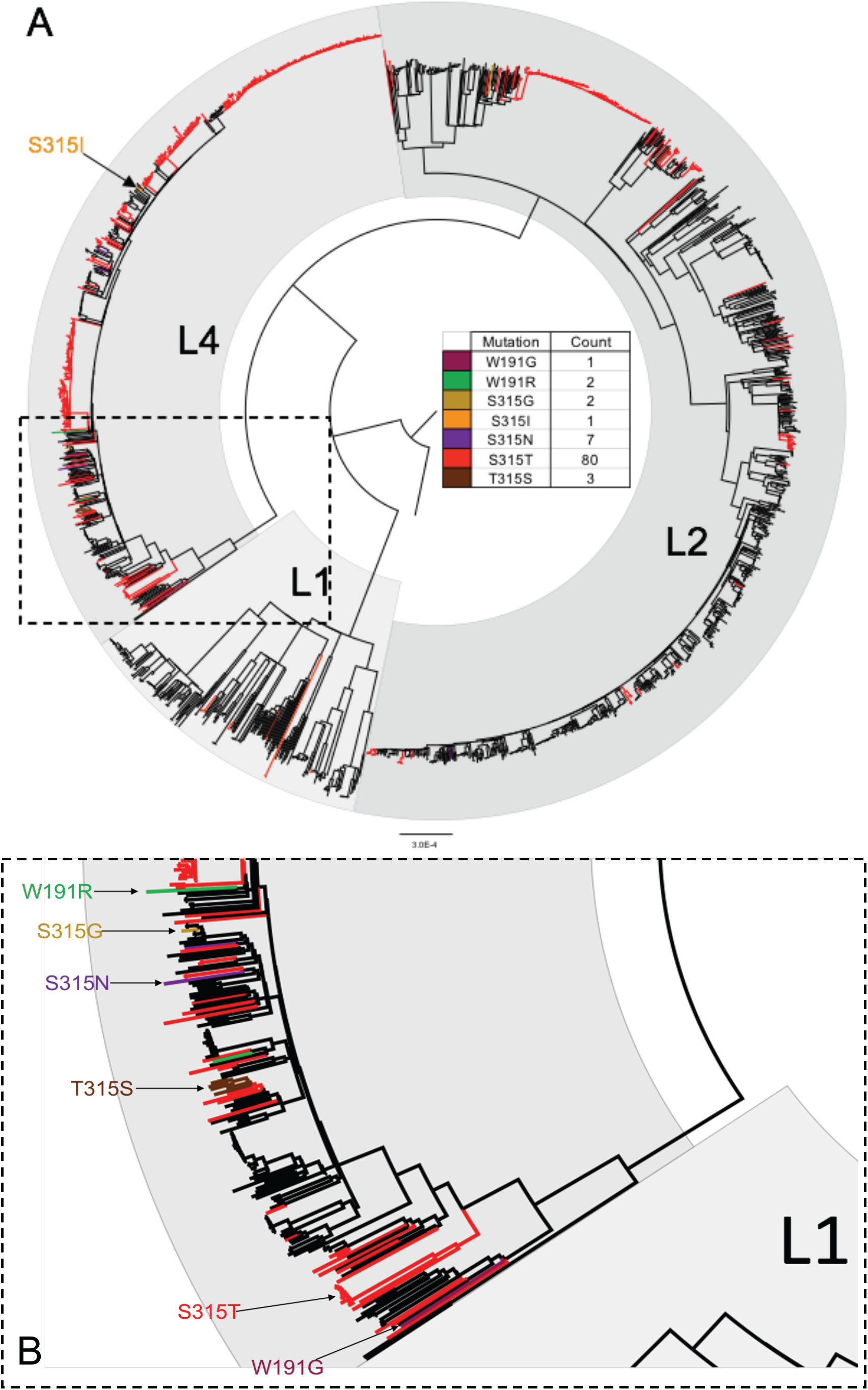
Real data example: *katG* homoplasies detected in *Mycobacterium tuberculosis*. (**A**) Maximum likelihood phylogeny showing location of homoplasic mutation events in the gene *katG* identified using SNPPar, coloured to indicate the resulting amino acid substitution (as per inset legend). (**B**) Expanded insert from **A**. Note that this analysis includes 2000 strains randomly sampled from the Global L124 set^31^; L1, L2, L4 indicate lineages 1, 2 and 4.

## 5. Discussion

Performance testing of SNPPar based on simulated data demonstrated that it has very high sensitivity to detect homoplasic mutation events (with 89% of tests yielding perfect recall, range 0-2 false-negatives per test) and also has high specificity (*i*.*e*. no false-positive calls were produced). The number of false-negative calls did increase slightly with increasing sample complexity (see **Figure 1**), hence the raw number of false-negatives may increase for larger empirical data sets. SNPPar was also highly accurate in classifying the identified homoplasies into types; the percentage of incorrect calls was inversely proportional to the sample size (see **Figure S2A**), with <1% of incorrect calls in the largest simulated dataset. Most homoplasy type-call errors were either revertant mutation events called as parallel events, or *vice versa*, where the call involved the root node and thus cannot be resolved by any method without the use of outgroups, or additional information about ancestral states (see **Figure S2B**).

Examination of the computational efficiency of SNPPar demonstrated a linear relationship between execution time and total alignment length for simulated data, when using the default TreeTime for ASR (see **Figure 5A**). Real data took more time and memory to execute due to a greater number of allele patterns to test, mainly due to the presence of missing alleles in real data. Of the three sorting methods tested, intermediate sorting was the quickest for both simulated and empirical datasets (see **Figure 5A**). However, this came with an increase in memory usage compared to the slowest, complex sorting, particularly with the larger datasets (see **Figure 5B**), as more SNPs are passed to TreeTime for ASR. Notably all three sorting methods were much more efficient than using TreeTime to analyse the entire alignment (∼4 GB for SNPPar with intermediate sorting and the largest empirical subsampled dataset *versus* ∼24 GB for TreeTime). As the time saving was notably higher for intermediate sorting, and memory usage was still in a reasonable range, we have set this sorting method as the default. If memory becomes limiting, *e*.*g*. on much larger data sets, complex sorting may be preferable but can be expected to take about twice as long. Considering where improvements in performance could be made in future for tackling much larger datasets, possible strategies include doing simple sorting to save time but farming out the ASR step into small batches to reduce memory, or first dividing the task before the sorting step (whilst ensuring SNPs in same gene stay within the same batch). Using FastML for the ASR step in SNPPar was notably slower for both simulated and real data, with no improvement in accuracy or memory use; hence we recommend using the default TreeTime for ASR.

SNPPar analysis made it relatively easy to reproduce the selection analyses reported previously on empirical data; one command, run using standard input files of tree and SNP alignment, took a few seconds on a laptop, producing details of all SNPs annotated with their coding effects and position in the tree. The richness of the output makes it straightforward to not only identify homoplasic mutation events, but also to explore convergent evolution events in detail, including by summarising the data at the levels of SNP site, codon and gene (see **Tables 1-2**) and visualising convergent and/or diversifying evolutionary across the genome and in the context of the phylogeny (see **Figures 6-9**). Notably, the SNPPar analysis is more reproducible than the published analyses and can be readily repeated as more isolates are added to the dataset.

There are some caveats with using SNPPar. Firstly, as with any algorithm, results from SNPPar are only as good as the input data that is provided. Prior quality control of the SNP alignment is important, not only to obtain high-confidence SNP allele calls by removing those found in repeat regions or other regions with ambiguous or low-quality calls (*e*.*g*. the (P)PE genes found in *Mtb*^31^), but also to improve the resulting tree by removing the noise from these potentially ambiguous sites. The phylogenetic tree itself must then be inferred from the data, and there may be multiple trees that describe the data equally well; indeed, a bifurcating tree may not capture the relationships between strains accurately, and there may be low confidence regions of the tree topology. All these factors should be considered when feeding input to SNPPar, and when interpreting the results. For example, where multiple trees describe the data equally well (e.g. where there are many polytomies) or any other phylogenetic uncertainty, each tree from a set of candidates (e.g. those with the same statistical fit or from bootstrap replicates) could be run through SNPPar and the results combined to obtain consensus mutation event data from which to infer selection. Secondly, SNPs can arise through spontaneous mutations (independent substitution events) or through recombination events (horizontal transfer of a group of linked variants); the latter need to be efficiently removed from the input SNP alignment if a user wishes to interpret SNPPar mutation event data as unlinked substitutions signifying independent parallel evolution. However, the output of SNPPar could be used to check an alignment for evidence of clusters of SNPs mapped to the same branch of the input tree, which may represent recombination; these could then be properly identified and removed from the alignment with targeted software (*e*.*g*. Gubbins^7^ or ClonalFrameML^8^) prior to re-running SNPPar with a recombination-filtered tree and SNP alignment. Finally, users should keep in mind the inherent limitation that ASR cannot accurately distinguish mutation events immediately below the root node^43,44^, and hence ensure to include an outgroup.

We have demonstrated that SNPPar works efficiently on large genome-wide SNP sets; it is highly accurate in detecting true positive homoplasic SNPs relative to an input tree with no false positive calls. SNPPar located homoplasic SNPs efficiently and accurately with both the simulated and real data sets tested here. Currently, SNPPar is the only tool that automates the detection and annotation of homoplasic SNPs efficiently from large SNP alignments. As further demonstrated by the real examples, this information can be used to explore the role of homoplasy in parallel and/or convergent evolution at the codon- and gene-level, in addition to identification of homoplasic SNPs *per se*. Whilst SNPPar was designed and tested specifically for the detection of homoplasic SNPs in bacteria, it could potentially be used for investigating homoplasic SNPs in other haploid organisms and viruses.

## Author statements

### Authors and contributors

*Conceptualisation*: KEH, DJE, BP, SD; *Methodology*: KEH, DJE, BP, SD; *Software*: DJE; *Validation*: DJE; *Formal Analysis*: DJE, KEH, BP, SD; *Resources*: KEH; *Data Curation*: DJE; *Writing (Original)*: DJE, KEH; *Writing (Review)*: DJE, KEH, BP,SD; *Visualisation*: DJE; *Supervision*: KEH, BP, SD; *Project Administration*: KEH, DJE; *Funding*: KEH.

## Acknowledgements

The authors would like to thank Dr Tami Lieberman for kindly providing the published version of the *B. dolosa* tree.

## Data bibliography

1. Holt KE, McAdam P, Thai PVK, Thuong NTT, Ha DTM, Lan NN, Lan NH, Nhu NTQ, Hai HT, Ha VTN, Thwaites G, Edwards DJ, Nath AP, Pham K, Ascher DB, Farrar J, Khor CC, Teo YY, Inouye M, Caws M, Dunstan SJ. Frequent transmission of the *Mycobacterium tuberculosis* Beijing lineage and positive selection for the EsxW Beijing variant in Vietnam. *Nat Genet* 2018;50:849-856 doi: 10.1038/s41588-018-0117-9
2. Perrin A, Larsonneur E, Nicholson AC, Edwards DJ, Gundlach KM, Whitney AM, Gulvik CA, Bell ME, Rendueles O, Cury J, Hugon P, Clermont D, Enouf V, Loparev V, Juieng P, Monson T, Warshauer D, Elbadawi LI, Spalding Walters M, Crist MB, Noble-Wang J, Borlaug G, Rocha EPC, Criscuolo A, Touchon M, Davis JP, Holt KE, McQuiston JR, Brisse S. Evolutionary dynamics and genomic features of the *Elizabethkingia anophelis* 2015 to 2016 Wisconsin outbreak strain. *Nat Commun* 2017;8:15483 doi: 10.1038/ncomms15483
3. Lieberman TD, Michel JB, Aingaran M, Potter-Bynoe G, Roux D, Davis MR Jr, Skurnik D, Leiby N, LiPuma JJ, Goldberg JB, McAdam AJ, Priebe GP, Kishony R Parallel bacterial evolution within multiple patients identifies candidate pathogenicity genes. *Nat Genet* 2011;43:1275-1280 doi: 10.1038/ng.997

