## Supplementary Information for "SNPPar: identifying convergent evolution and other homoplasies from microbial whole-genome alignments"

### *Internal logic for sorting non-singleton bi-allelic SNPs into monophyletic or paraphyletic within SNPPar*

At SNP sites with missing base calls, tips with unknown base calls are removed from the tree against which monophyly is then assessed (**Figure 3B**). If only one allele is monophyletic on the tree then the mutation event can be unambiguously assigned to the branch leading to this monophyletic group (**Figure 3C**), however if both alleles are monophyletic then the placement of the event relative to the tree root is ambiguous and the event is assigned arbitrarily to the branch leading to the minor (i.e. less frequent) allele (**Figure 3D**). Assessment of monophyly is conducted on allele patterns (*i.e.* for each SNP, the unique set of input sequences with the minor, major, and missing alleles respectively) and the results are cached, to reduce repeat computation for the allele patterns that occur multiple times in the data set.

### *Annotation of the coding effects of mutation events mapped to the tree within SNPPar*

Coding domain sequences (CDS) are labelled using the locus_tag qualifier (if these qualifiers are not present in the reference genome, they are created on the fly). Intergenic SNPs are annotated with the distance to, and orientation of, the nearest CDS in each direction. Intragenic SNPs are annotated with the CDS in which they fall, along with the codon coordinate within that CDS, the ancestral and derived codon sequences and amino acid residues, and an indicator of the amino acid change (*i.e.* synonymous, nonsynonymous or ambiguous). To ensure that the coding effect of point mutations in CDS are calculated accurately, it is necessary to consider codon triplets rather than individual nucleotides, in the specific context of the branch to which the mutation event for a SNP has been mapped. SNPPar achieves this by first collating internal node sequences for all sites (combining those output by TreeTime with those inferred internally), then assessing the effect of a given mutation event mapped to a given branch by considering the change in the triplet codon from the parent node to the child node. For SNP sites that lie in multiple overlapping protein-coding genes, each SNP effect is reported on a new line (note that unique mutation events can be trivially retrieved from this coding effect list as the unique combination of site, branch and base substitution).

### *Simulation of* Mtb *sequence data for assessing accuracy of SNPPar homoplasy calls*

SeqGen v1.3.4[1] was used to simulate DNA sequence evolution on each empirical tree, utilising a GTR+G model with the corresponding parameter values estimated from each of the 12 empirical datasets. The starting sequence was a version of the *Mtb* reference genome, H37Rv, shortened to remove the total length of sites that were filtered out when extracting core SNPs for the original *Mtb* empirical data analysis[2]. Following an initial simulation run with each data set, branch lengths in the input trees were scaled to ensure the branch lengths correspond to substitutions per SNP site (as per the empirical data) rather than substitutions per site (which includes

Company Limited by Guarantee l Registered in England No. 1039582 l Registered Office Charles Darwin House, 12 Roger Street, WC1N 2JU, UK
Registered as a Charity: 264017 (England & Wales); SC039250 (Scotland)

sites with no polymorphisms). SeqGen returns simulated sequences at each node in the tree (including tips) for the complete input sequence. We therefore used Python scripts (available in the repository) to first extract the variable sites, then traverse the input tree and compare simulated parent-child node sequence pairs, in order to identify mutation events on each branch and thus identify homoplasic SNPs (and their exact type). A SNP allele table was extracted from each set of simulated tip sequences to use as input for SNPPar.

#### *Analysis of Variance (ANOVA) of SNPPar performance based on simulated data*

**Model:** Time ~ Length * Sorting
(where Length = total alignment length, i.e. number of SNP sites multiplied by number of samples; Sorting = categorical variable with values simple, intermediate, complex)

| ANOVA | | | | | |
|---|---|---|---|---|---|
| **Response:** Time | **Df** | **Sum Sq** | **Mean Sq** | **F value** | **Pr(>F)** |
| Length | 1 | 12323.4 | 12323.4 | 93660.14 | < 2.2e-16 *** |
| Sorting | 1 | 14.4 | 14.4 | 109.67 | < 2.2e-16 *** |
| Length:Sorting | 1 | 50.1 | 50.1 | 380.41 | < 2.2e-16 *** |
| Residuals | 236 | 31.1 | 0.1 | | |

**Model:** Memory ~ SNPs * Sorting
(where SNPs = number of SNP sites; Sorting = categorical variable with values simple, intermediate, complex)

| ANOVA | | | | | |
|---|---|---|---|---|---|
| **Response:** Memory | **Df** | **Sum Sq** | **Mean Sq** | **F value** | **Pr(>F)** |
| SNPs | 1 | 305099696 | 305099696 | 3339.42 | < 2.2e-16 *** |
| Sorting | 1 | 35480904 | 35480904 | 388.35 | < 2.2e-16 *** |
| SNPs:Sorting | 1 | 46081936 | 46081936 | 504.38 | < 2.2e-16 *** |
| Residuals | 236 | 21561680 | 91363 | | |

#### *References*


1. Rambaut A, Grass NC. Seq-Gen: an application for the Monte Carlo simulation of DNA sequence evolution along phylogenetic trees. *Bioinformatics* 1997;13:235–238 doi: 10.1093/bioinformatics/13.3.235

2. Holt KE, McAdam P, Thai PVK, Thuong NTT, Ha DTM, Lan NN, Lan NH, Nhu NTQ, Hai HT, Ha VTN, Thwaites G, Edwards DJ, Nath AP, Pham K, Ascher DB, Farrar J, Khor CC, Teo YY, Inouye M, Caws M, Dunstan SJ. Frequent transmission of the *Mycobacterium tuberculosis* Beijing lineage and positive selection for the EsxW Beijing variant in Vietnam. *Nat Genet* 2018;50:849-856 doi: 10.1038/s41588-018-0117-9

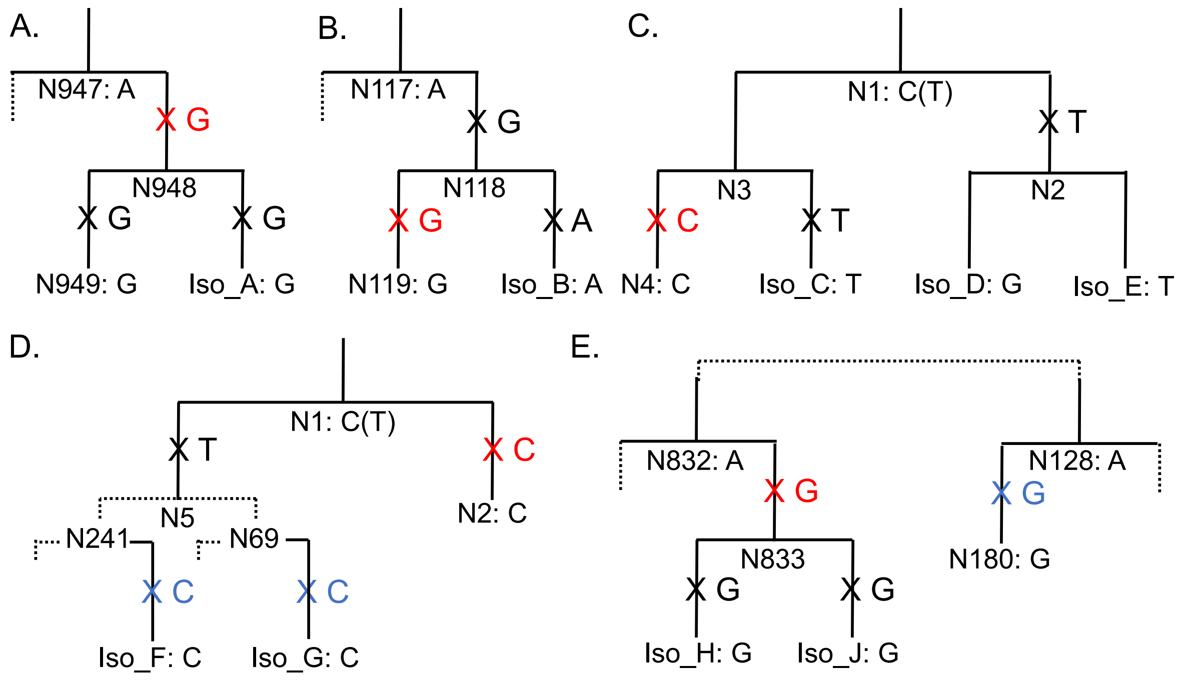


**Supplementary Figure S1. Examples of false-negative homoplasy calls (A-C) and incorrect type calls (D-E) made by SNPPar when analysing simulated *Mtb* data.** Crosses indicate mutation events, black crosses indicate simulated mutations that were missed, red crosses indicate those called incorrectly by SNPPar, blue crosses indicate those correctly called by SNPPar. Branch labels starting with 'N' are internal nodes (N1 being the root node of the tree), tips are labelled as Iso_'X'. The letter after the colon indicates the nucleotide present at each node; for the root node (i.e. N1), the incorrectly (arbitrarily) inferred nucleotide is also shown in brackets. Branch lengths are not to scale. **A.** False-negative call at position 2,486,385 in Global L124, 2,000 isolates, run 5 (of 10). Expected: parallel mutation event, called: single mutation event. **B.** False-negative call at position 165,037 in Global L124, 2,000 isolates, run 9. Expected: revertant mutation event, called: single mutation event. **C.** False-negative call at position 2,499,720 in Global L2, 10% of isolates, run 4. Expected: parallel mutation event, called: single mutation event. **D.** Incorrect type call at position 3,562,937 in Global L2, all isolates, run 3. Expected: revertant mutation event in parallel, called: parallel mutation event. **E.** Incorrect type call at position 3,755,767 in Global L124, 1,000 isolates, run 3. Expected: parallel mutation events (by three), called: parallel mutation event (by two, i.e. one is missed).

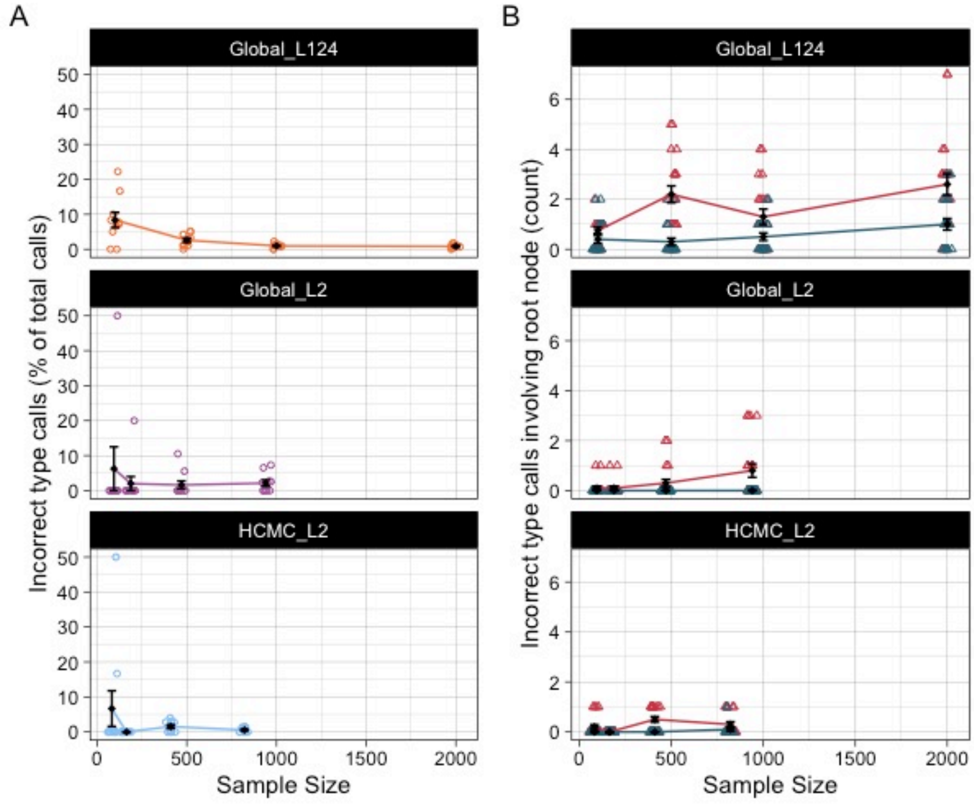


**Supplementary Figure S2. Incorrect homoplasy type calls for the three populations found using the simulated datasets and TreeTime: A. Percentage of incorrect homoplasy type calls for all true positive (correct) homoplasy calls; B. Number of incorrect type calls for homoplasies involving only two mutation events (*i.e.* parallel and reversion events) where the root node is involved.** For each sample size, n=10 replicates. In **B.**, red triangles indicate expected reversions called as parallel, whilst blue triangles indicate expected parallel events called as reversions. Means and standard errors are indicated by the black points and crossbars. The six incorrect type calls included in **A.** but excluded from **B.** either don't involve the root node (four type calls: three reversions called as parallel and one parallel event called as a reversion) or involved three mutation events (two type calls, one involving a root node call where a parallel reversion was instead called as three parallel events and the other (*sans* root node) where two parallel events of the expected three were collapsed into a single event as they occurred on sister branches, see **Figures S1D-E** for examples).

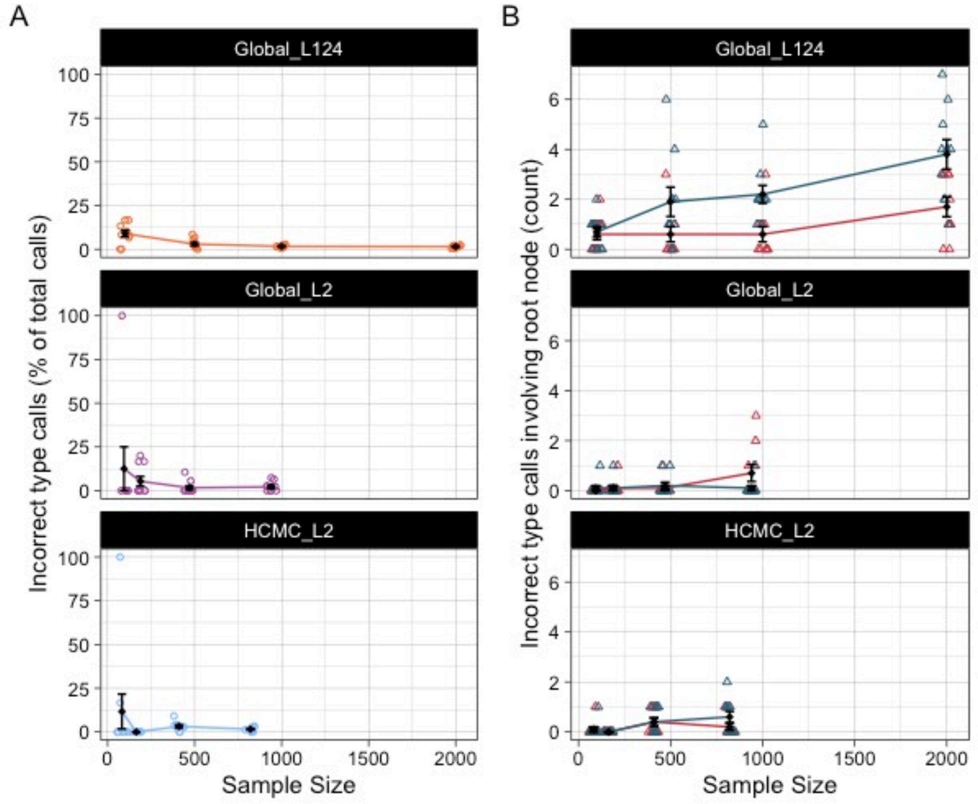


**Supplementary Figure S3. Incorrect homoplasy type calls for the three populations found using the simulated datasets and FastML: A. Percentage of incorrect homoplasy type calls for all true positive (correct) homoplasy calls; B. Number of incorrect type calls for homoplasies involving only two mutation events (*i.e.* parallel and reversion events) where the root node is involved.** For each sample size, n=10 replicates. In **B.**, red triangles indicate expected reversions called as parallel, whilst blue triangles indicate expected parallel events called as reversions. Means and standard errors are indicated by the black points and crossbars.

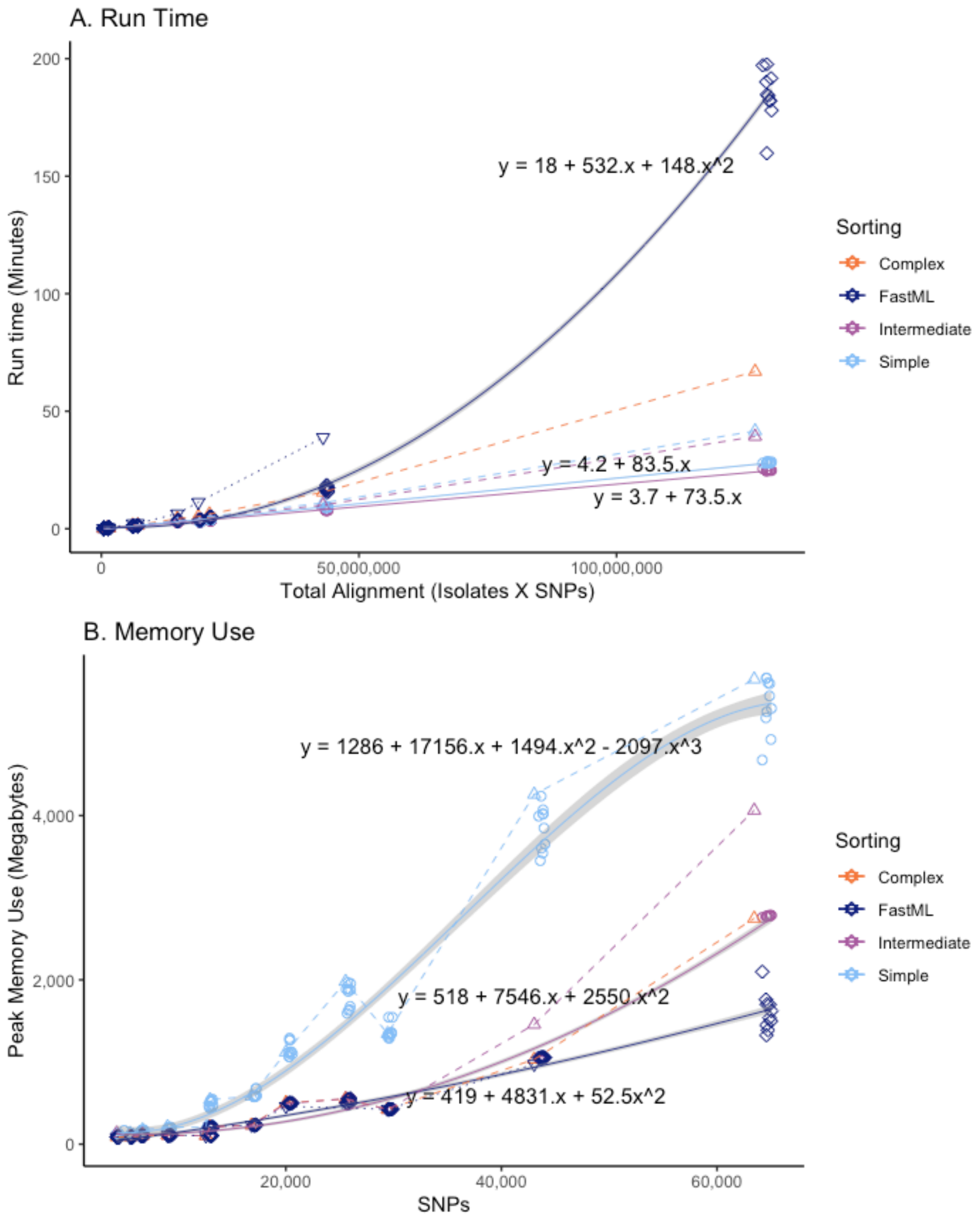


**Supplementary Figure S4. SNPPar performance during analysis of subsampled empirical and simulated datasets using different sorting options, including comparing FastML to TreeTime for ASR.** (**A**) Total run time vs total alignment length. (**B**) Peak memory use vs SNP alignment length. Circles denote simulated datasets, triangles empirical datasets; colours indicate sorting options as per inset legend. The simulated datasets have ten replicates of each sample size, and a line of best fit through these is indicated (solid line, grey shading indicates 90% fit interval). Note complex sorting was not used for the simulated datasets as there are no missing calls, and complex and intermediate sorting are identical in this case. The real datasets have only a single observations per sorting algorithm, which are joined with simple dashed lines.
